# The neural correlates of two forms of spiritual love: an EEG study

**DOI:** 10.1101/045898

**Authors:** E. S. Louise Faber

## Abstract

Spiritual practices are gaining an increasingly wider audience as a means to enhance positive affect in healthy individuals and to treat neurological disorders such as anxiety and depression. The current study aimed to examine the neural correlates of two different forms of love generated by spiritual practices using EEG; love generated during a loving kindness meditation performed by Buddhist meditators, and love generated during prayer, in a separate group of participants from a Christian-based faith. The loving kindness meditation was associated with significant increases in delta, alpha 1, alpha 2 and beta power compared to baseline, while prayer induced significant increases in power of alpha 1 and gamma oscillations, together with an increase in the gamma: theta ratio. An increase in delta activity occurred during the loving kindness meditation but not during prayer. In contrast increases in theta, alpha 1, alpha 2, beta and gamma power were observed when comparing both types of practice to baseline, suggesting that increases in these frequency bands are the neural correlates of spiritual love, independent of the type of practice used to attain the state of this type of love. These findings show that both spiritual love practices are associated with widespread changes in neural activity across the brain, in particular at frequency ranges that have been implicated in positive emotional experience, integration of distributed neural activity, and changes in short-term and longterm neural circuitry.

## Introduction

In recent years there has been a growing interest within cognitive neuroscience to better understand the neural mechanisms of spiritual practices, especially different forms of meditation, due to its widely reported beneficial effects (Goyal *et al*., 2014). In addition to being used in non-clinical settings to alleviate stress and induce positive affect, these practices have also been used in clinical settings as complimentary treatments for psychiatric disorders, such as anxiety and depression (Baer, 2003; Grossman *et al*., 2004; Sundquist *et al*., 2014). The aim of the current study was to examine, using electroencephalography (EEG), the neural correlates of love generated with spiritual practices. “Spiritual love” differs from maternal love or romantic love since it is generated through spiritual practices such as meditation or prayer. Two different methods for generating spiritual love were examined; a non-theistic love state acquired through the Buddhist loving kindness meditation (LKM), which involves the generation of non-referential love and compassion, and a theistic love state acquired through prayer, during which participants from a Christian-based faith cultivate a feeling of love towards God, and desire to receive love from God.

A number of studies have examined the effects of the experience of different forms of love on neural activity. Romantic love is associated with activation of cortical regions involved in affective processing (Davidson & Irwin, 1999), such as the medial insula and the anterior cingulate cortex (Bartels & Zeki, 2000), in addition to subcortical regions involved in reward circuitry (Kelley & Berridge, 2002), such as the ventral tegmental area, caudate nucleus and putamen (Bartels & Zeki, 2000; Aron *et al*., 2005; Xu *et al*., 2011). Similarly, maternal love involves activation of the middle insula and the dorsal anterior cingulate cortex, as well as deactivation in the amygdala (Bartels & Zeki, 2004). Unconditional love is also correlated with activation of subcortical reward centres, including the left ventral tegmental area and right caudate nucleus, and regions involved in affect, such as the left anterior cingulate cortex and the insula (Beauregard *et al*., 2009). Only one EEG study has examined neuronal oscillations associated with romantic love, reporting significant increases in the delta frequency power during the experience of romantic love (Basar *et al*., 2008).

By contrasting the practices of a LKM and prayer, we sought to elucidate the neural correlates that are common to both forms of spiritual love and therefore not solely attributable to the cognitive activity used to acquire that state of spiritual love. A previous study investigating the neural correlates of spiritual love using EEG, in which the LKM was examined (Lutz *et al*., 2004), found increases in gamma power as both a state and trait effect in experienced Tibetan Buddhist practitioners. In the current study we compared the electrophysiological responses of moderately experienced meditators performing a Tibetan Buddhist LKM with a separate group of participants carrying out prayer.

## Methods

### Participants

Twenty-eight participants of Caucasian origin were recruited for this study. Sixteen (12 females, 4 males) were moderately experienced Buddhist meditators (mean age = 43.3 years, SD = 11.6, range: 25-58 years) and twelve (7 females, 5 males) were moderately experienced practitioners of “Divine Truth” (mean age = 37.4 years, SD = 8.4 years, range: 25-47 years), a Christian-based spiritual practice that teaches prayer towards God, involving cultivating a feeling of love towards God and a desire to receive love from God. There was no significant difference between the ages of the two groups of participants (Student's *t* test; *p* > 0.05), nor any difference in gender or handedness (Fisher's Exact Probability test; *p* > 0.05). Meditators had trained in Tibetan Buddhist meditation for between 800-3800 hours, with an average of 1,900 hours practice, relating to an average of 45 minutes practice per day for 7 years. Prayer participants had carried out between 500-8000 hours of practice, with an average of 1,700 hours of practice, over a period of 0.8 to 6 years. Practitioners of both groups incorporated the spiritual practices into their daily routine around education and employment commitments. The study was approved by the University of Queensland Human Research Ethics Committee. Experiments were undertaken with the understanding and written consent of each participant, and the study conforms with World Medical Association Declaration of Helsinki. Participants were compensated $20 for the 2 hour study. All participants had no history of any medical disorder or neurological illness, and were not currently on any psychoactive drugs. Participants were blinded to specific experimental hypotheses.

### EEG recording and protocol

Continuous EEG data were recorded at a sampling frequency of 1024 Hz from 64 active scalp Ag-AgCl electrodes placed in an elastic cap, arranged according to the international standard 10-10 system (Oostenveld & Praamstra, 2001), using the BioSemi Active Two system (BioSemi, Amsterdam, Netherlands). Electro-oculogram (EOG) was recorded by placing electrodes lateral to the outer canthus of each eye and above and below the left eye, in line with the pupil, to monitor horizontal and vertical eye movements respectively. Electromyographic (EMG) activity was also recorded by placing electrodes over the right corrugator supercilii muscle (for frowning movements) and right zygomaticus major muscle (for smiling movements). However, these data will not be presented here. The electrodes were fixed using a gel and electrode offsets were kept below 30 *μ*V to ensure optimal connectivity before the start of each recording. During data acquisition, the EEG signal was referenced to the CMS (Common Mode Sense active) and DRL (Driven Right Leg passive) electrodes. All recordings were performed in a Faraday sealed room, to eliminate external sources of electrical artifacts, at the Queensland Brain Institute at the University of Queensland.

### Experimental procedure

Participants were instructed to avoid caffeine and nicotine on the day of the recording, and were asked to arrive well rested, in order to minimize the known effects of these facets on neuronal oscillations (Niedermeyer & Lopez da Silva, 1993). Participants were briefed about the experiment and filled in the consent form and two questionnaires, the Mindfulness and Attention Awareness scale (Brown & Ryan, 2003) and the Intrinsic Spirituality Scale (Hodge, 2003). The results of those questionnaires will not be presented in the current report.

For resting EEG, four blocks of 60 seconds were recorded, two with eyes open and two with eyes closed, in a counterbalanced order. During this baseline period, participants were instructed to be in a neutral, relaxed state of mind and were asked to focus on a fixation cross that was presented on the computer screen. Meditators underwent four blocks each of two different practices in a pseudorandomised order, the LKM and a focused attention meditation. Similarly, prayer participants underwent four blocks each of two different practices presented in a pseudo-randomised order. The first involved the participants praying towards God by asking God for love and cultivating a feeling of love towards God, cued by the term "Divine Love" on the computer screen. The second practice asked participants to think about God, but not to engage in a prayer. Only the practices that generated spiritual love (i.e. LKM for the Buddhist practitioners and prayer for the Divine Truth participants) are analyzed and discussed here.

Each block involved a 30 second baseline period instructing the participant to get ready. This was followed by a 70 second session with their eyes open, where the meditators performed the LKM or a focused attention meditation, and the prayer practitioners carried out prayer or the contemplating God practice. The time lapsed during this session was indicated by a timer on the computer screen, which the participants were asked to fixate upon. Both groups of participants indicated the onset and offset of their respective practices by a button press, to enable precise extraction of the data involving the spiritual love state. During the prayer practice, participants indicated the onset button press when they felt they were experiencing a loving connection with God. The end of the practice was followed by a 30 second rest period (Figure 1). A three point rating scale was then presented on the computer screen, to which the participants were asked to rate the quality of the practice, with 1 indicating a poor quality block, 2 indicating a mediocre quality block and 3 indicating a high quality block. Only the top 3 rated practice blocks for each participant were used for further analysis.

**Figure 1.**
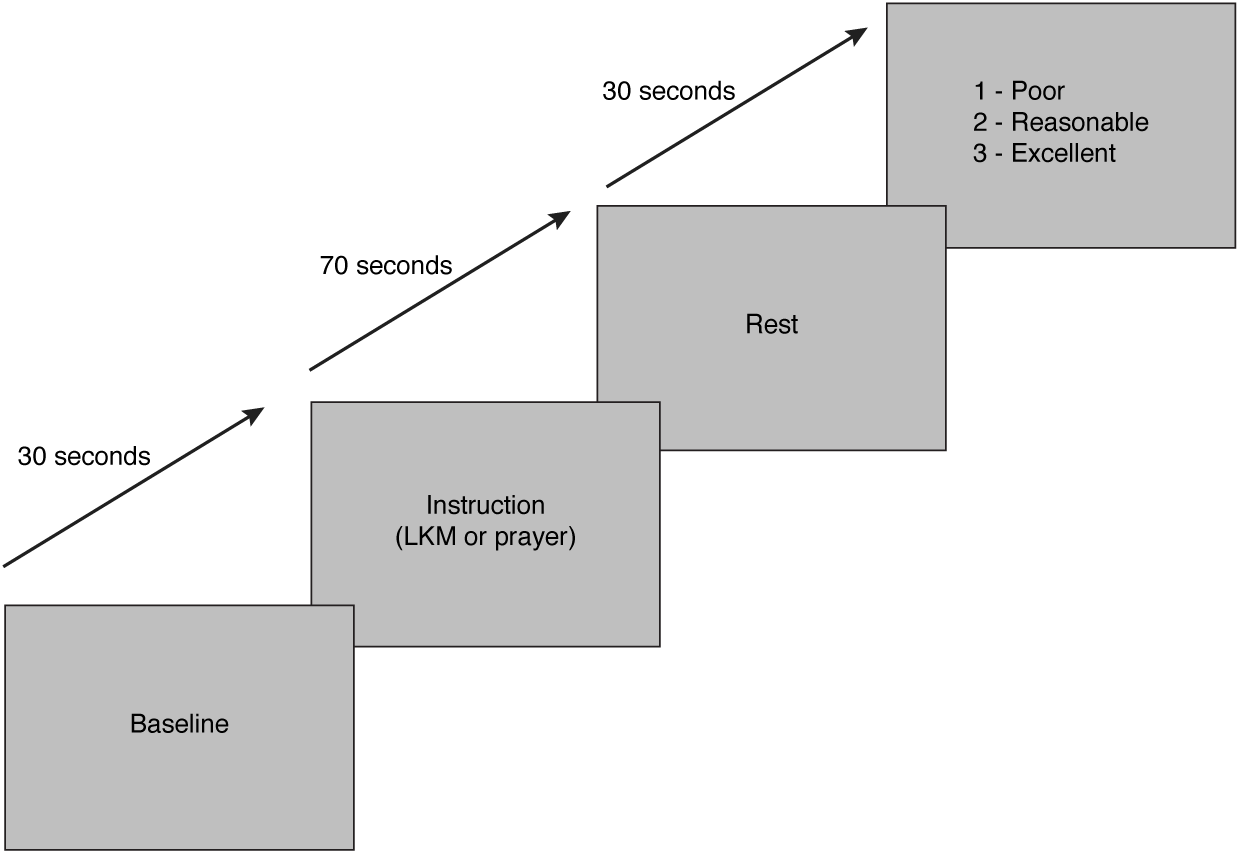
Experimental paradigm. Schematic figure of a single block (out of 8) of instructions, with the respective time intervals, provided to participants on screen. Participants were first presented with a "Baseline" instruction, lasting 30 seconds. Next followed a 70 second session, during which meditators were presented with the instruction “Loving Kindness Meditation”, whereas prayer participants were presented with the instruction “Divine Love”. This was followed by a short “Rest” block of 30 seconds, ending with a rating scale to assess the quality of the practice session.

### EEG spectral analysis

For each participant, raw EEG data was re-referenced offline to the average of all the scalp channels and then digitally filtered using a zero phase shift high pass filter at 0.1 Hz with a 6db/octave. Pre-processing involved rejection of EEG sections with muscular artifacts and removal of EOG artifacts using BESA v5.3 (MEGIS Software, Inc., Gräfelfing, Munich). One meditator’s data could not be processed further due to high levels of noise in their EEG trace and was excluded from statistical analysis. The continuous data were then imported into Neuroscan Edit 4.4 (Compumedics Inc., Charlotte, NC) and subjected to a zero phase shift low-pass filter at 45 Hz with a 48dB/octave. For each condition of interest, data were segmented into non-overlapping two second epochs and baseline corrected across the entire sweep. Using an automatic artifact rejection process, epochs in which amplitude exceeded ± 75μV were rejected. Epochs still containing any remnant muscular, ocular or other artifacts were visually inspected and removed. The resulting epoched data were then averaged in the frequency domain using a fast Fourier Transform (FFT) with a Hanning window (10% taper). The EEG power spectrum was then examined in the following bands: delta (2-4 Hz), theta (4-8 Hz), alpha 1 (8-10 Hz), alpha 2 (10-13 Hz), beta (13-30 Hz) and gamma (30-44 Hz). The alpha band (8-13 Hz) was subdivided into alpha 1 and alpha 2 since they are known to have different functional roles (Klimesch *et al*., 1999). Inspection of histograms and normality probability plots indicated that power values (μV^2^) in each frequency band were not normally distributed and hence were log-transformed. The ratios of the gamma frequency to slower frequency bands were also examined since slow oscillations (4-13 Hz) are known to play a complimentary role to the faster rhythms (Ward, 2003), and have been implicated in meditative practices generating love (Lutz *et al*., 2004). Gamma: theta, gamma: alpha 1 and gamma: alpha 2 values were calculated individually rather than grouping them collectively as slow oscillations (4-13 Hz) in order to pinpoint which of the slow oscillation(s) play a significant complementary role.

For the purposes of statistical analyses, the power and gamma ratio data from 21 electrodes (Fp1, Fpz, Fp2, F7, F3, Fz, F4, F8, T7, C3, Cz, C4, T8, P7, P3, Pz, P4, P8, O1, Oz and O2) were used to provide extensive scalp coverage. Using SPSS version 18 (SPSS Inc., Chicago, IL), each frequency band and gamma ratio was subjected to separate repeated measures analysis of variance designs (ANOVA) for meditators and DLP practitioners with factors including “electrode” (21 electrodes) and “state” (love vs. baseline). The love state showed power values from either the LKM or prayer, and the baseline state showed power values from the resting EEG during the initial eyes open condition, before any of the love practices had commenced, so as to avoid any inter-practice effects on the baseline data. Differences between the two spiritual practices were examined using a mixed design ANOVA with the aforementioned repeated measures of “electrode” and “state”, and a between groups factor of “group” (meditators vs. prayer practitioners). Huynh Feldt corrections were applied where sphericity was violated, as determined by a significant Mauchley’s test. Statistical significance was set at *p* < 0.05 for each of the tests with Bonferroni adjusted *p* values used to correct for multiple comparisons.

## Results

EEG signals were acquired to examine the effects of the two different spiritual love practices. Power spectrum analysis was carried out in order to gain a measure of increases in neuronal activity in certain frequency bands, which results from increased neuronal oscillations in a common phase (Niedermeyer, 1993 #227).

### Loving kindness meditation

During the LKM, significant increases in delta (2-4 Hz), alpha 1 (8-10 Hz), alpha 2 (10-13 Hz) and beta (13-30 Hz) frequency bands were observed compared to baseline (Figure 2). However no significant changes were found in the theta frequency band (*F*(20, 280) = 1.964, *p* = 0.128, 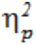 = 0.123, ε = 0.164) and, in contrast to a previous study (Lutz *et al*., 2004), no increases were observed in the gamma frequency band (*F*(20, 280) = 0.590, *p* = 0.752, 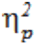 = 0.04, ε = 0.328).

**Figure 2.**
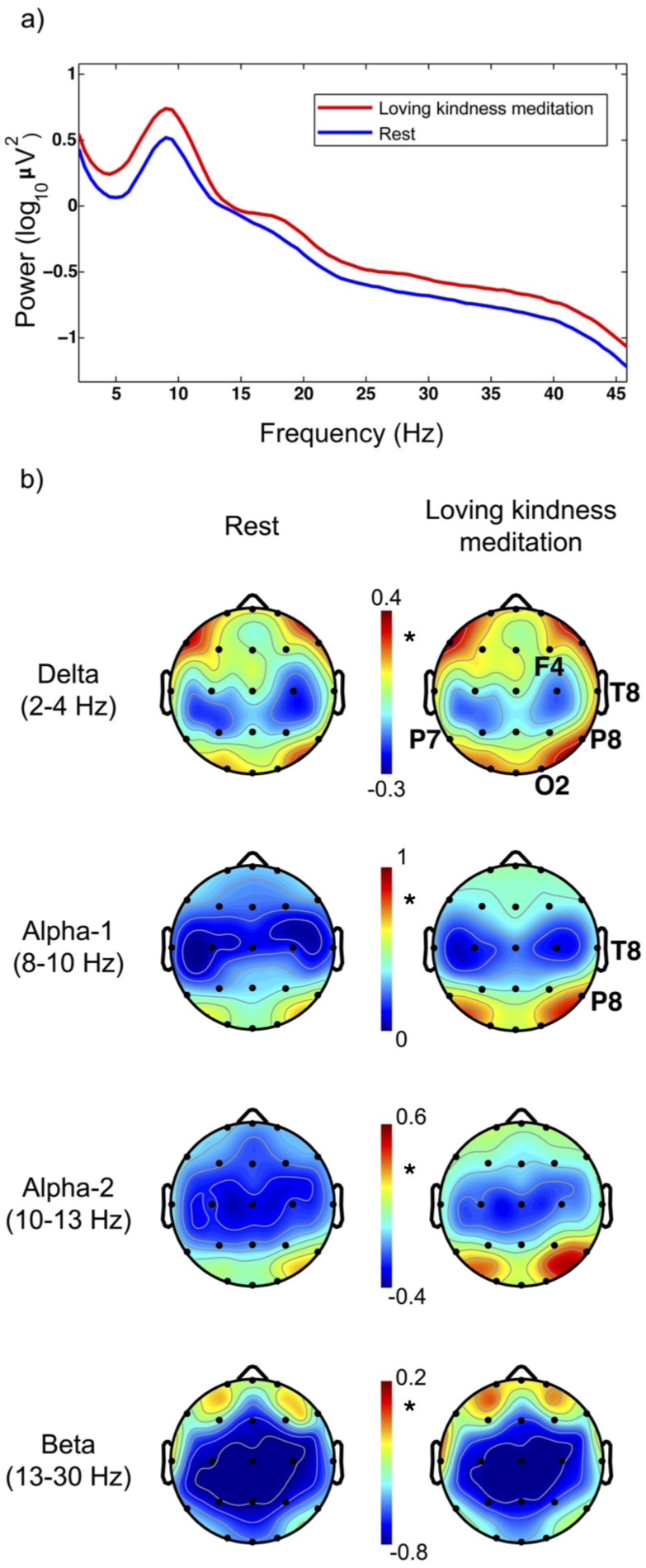
The neural correlates of the loving kindness meditation. a) The power spectrum for electrode P8 during the baseline condition (in blue) and the loving-kindness meditation (in red). b) Topographical maps for the delta, alpha 1, alpha 2 and beta bands are displayed during baseline conditions in the left panels and during the loving-kindness meditation (LKM) on the right. Statistical significance was determined using an ANOVA with multiple comparisons correction and significance set at *p*<0.05. Warmer colours denote greater activation. The labeling of an electrode indicates a significant increase seen at that individual electrode, while significant increases in power averaged across all electrodes are indicated by an asterisk next to the colour bar between the panels in this and subsequent figures. Significant increases in delta (F4, T8, P7, P8 and O2), alpha 1 (T8 and P8), alpha 2 and beta power were seen during the LKM compared to baseline.

A significant interaction effect between state and electrode was observed for the delta frequency band (*F*(20, 280) = 3.158, *p* < 0.001, 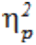 = 0.184, ε = 0.609; Figure 2). Bonferroni corrected pairwise comparisons revealed increases in delta power during the LKM compared to baseline at F4, T8, P7, P8 and O2 (*p* < 0.05 for all electrodes). Similarly, a significant interaction between state and electrode was observed in the alpha 1 frequency band (*F*(20, 280) = 2.857, *p* = 0.015, 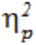 = 0.169, ε = 0.295; Figure 2), with pairwise comparisons showing significantly increased alpha 1 power during the LKM compared to baseline at right lateral temporal and parietal electrodes (T8 and P8, *p* < 0.05). For the alpha 2 frequency band, a main effect of state was observed with an increase in power observed during the LKM relative to baseline (*F*(1, 14) = 6.016, *p* = 0.028, 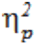 = 0.301; Figure 2). Similarly, a main effect of state was also observed for beta activity with significant increases observed during the LKM compared to baseline (*F*(1, 14) = 4.836, *p* = 0.045, 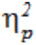 = 0.257; Figure 2).

No changes were observed in the ratio of gamma to theta (*F*(20, 280) = 1.130, *p* = 0.352, 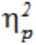 = 0.075, ε = 0.204), alpha 1 (*F*(20, 280) = 0.813, *p* = 0.501, 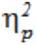 = 0.055, ε = 0.161) or alpha 2 frequency bands (*F*(20, 280) = 0.537, *p* = 0.656, 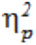 = 0.037, ε = 0.147).

### Prayer

In contrast to the LKM, during prayer significant increases in the alpha 1 and gamma frequency power were observed, in addition to an increase in the ratio of gamma to theta activity (Figure 3). Increases that showed a trend towards significance were also observed during prayer in the alpha 2 and beta frequency bands, whereas no significant changes were observed in the delta and theta frequency bands.

**Figure 3.**
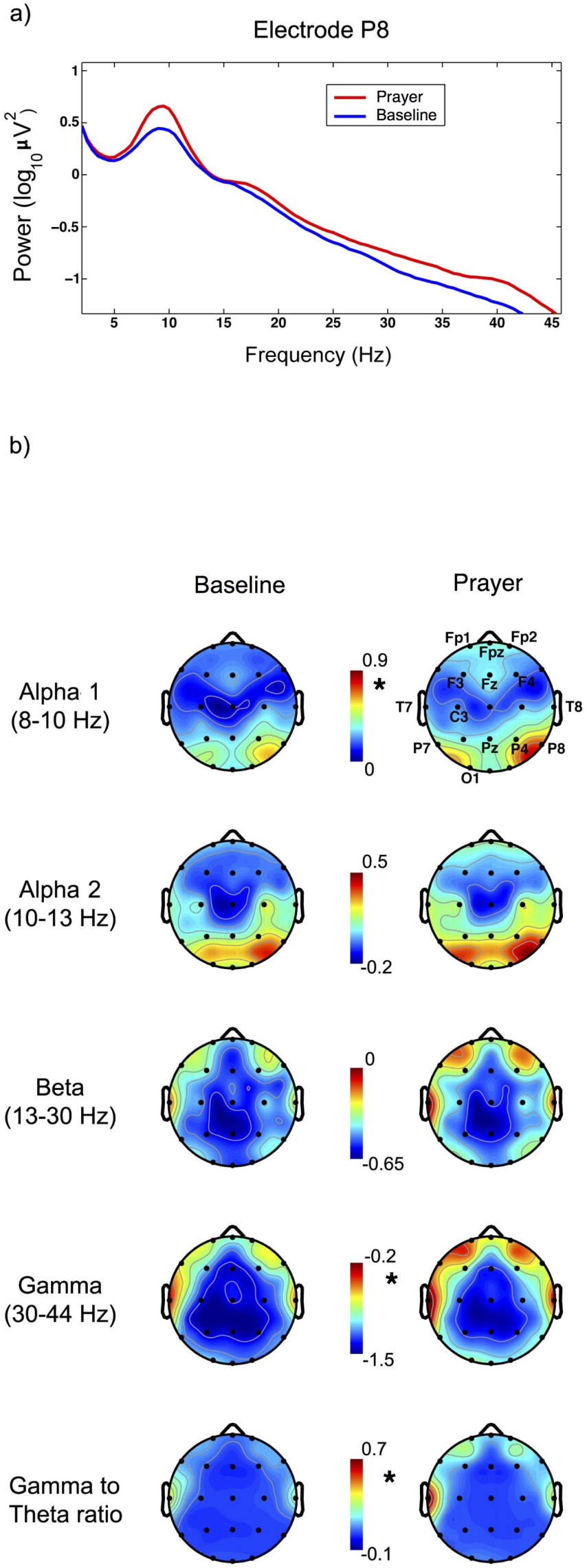
The neural correlates of prayer. a) The power spectrum for electrode P8 during the baseline condition (in blue) and prayer (in red). b) Topographical head maps for alpha 1, alpha 2, beta, gamma and gamma to theta are displayed on the left for the baseline condition and on the right for the prayer practice. Significantly increased power was observed in the alpha 1, gamma and gamma to theta ratio during prayer. A trend towards increased alpha 2 and beta power during prayer was also observed, however this did not reach statistical significance.

Post-hoc pairwise comparisons revealed that the significant increase in alpha 1 power during prayer (interaction effect between state and electrode; *F*(20, 220) = 2.733, *p* = 0.034, 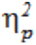 = 0.199, ε = 0.224; Figure 3) occurred over multiple electrodes, particularly over frontal electrodes, at Fp1, Fpz, Fp2, F3, Fz, F4, C3, T7, T8, Pz, P4, P7, P8 and O1 (*p* < 0.05 for all electrodes). Furthermore, a significant main effect of state was present for the gamma frequency band with increases observed during prayer relative to rest (*F*(1, 11) = 5.296, *p* = 0.042, 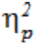 = 0.325; Figure 3). While no significant effects between state and electrode were observed for the alpha 2 frequency band, main effects revealed an increase in alpha 2 power during prayer compared to baseline that almost reached significance (*F*(1, 11) = 4.655, *p* = 0.054, 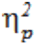 = 0.297; Figure 3). Similarly, a trend towards a significant increase in beta power was also observed during prayer compared to baseline (*F*(1, 11) = 4.587, *p* = 0.055, 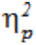 = 0.294; Figure 3).

Finally, the ratio of the gamma to theta frequency band, which is likely to play an important functional role coordinating activity across the brain (Varela *et al*., 2001), significantly increased during prayer relative to baseline as revealed by a main effect of state (*F*(1, 11) = 5.235, *p* = 0.043, 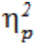 = 0.322; Figure 3). However no changes in the ratio of gamma to alpha 1 (*F*(20, 220) = 0.839, *p* = 0.418, 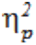 = 0.071, ε = 0.075) or alpha 2, (*F*(20, 220) = 0.956, *p* = 0.403, 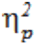 = 0.080, ε = 0.105) frequency bands were observed. This complementary increase in the ratio of fast (gamma) to slow (theta) frequency bands suggests a functional role of prayer in synchronizing fast neuronal activity in the gamma frequency band with slower theta range activity over large areas of the brain (Varela *et al*., 2001), and suggests a particular role for learning and memory (Rutishauser *et al*., 2010).

### Comparison between the spiritual practices

To examine any differences and similarities between prayer and the LKM, a mixed model ANOVA was used. For the delta frequency, a significant increase relative to baseline was observed only during the LKM and not during prayer, relative to baseline, as revealed by a significant three way interaction between state, electrode and group (*F*(20, 500) = 1.746, *p* = 0.049, 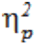 = 0.065, ε = 0.663; Figure 4a). Pairwise comparisons revealed that this increase in delta power during LKM was observed at F4, T8, P7, P8 and O2 (*p*<0.05 for all electrodes).

**Figure 4.**
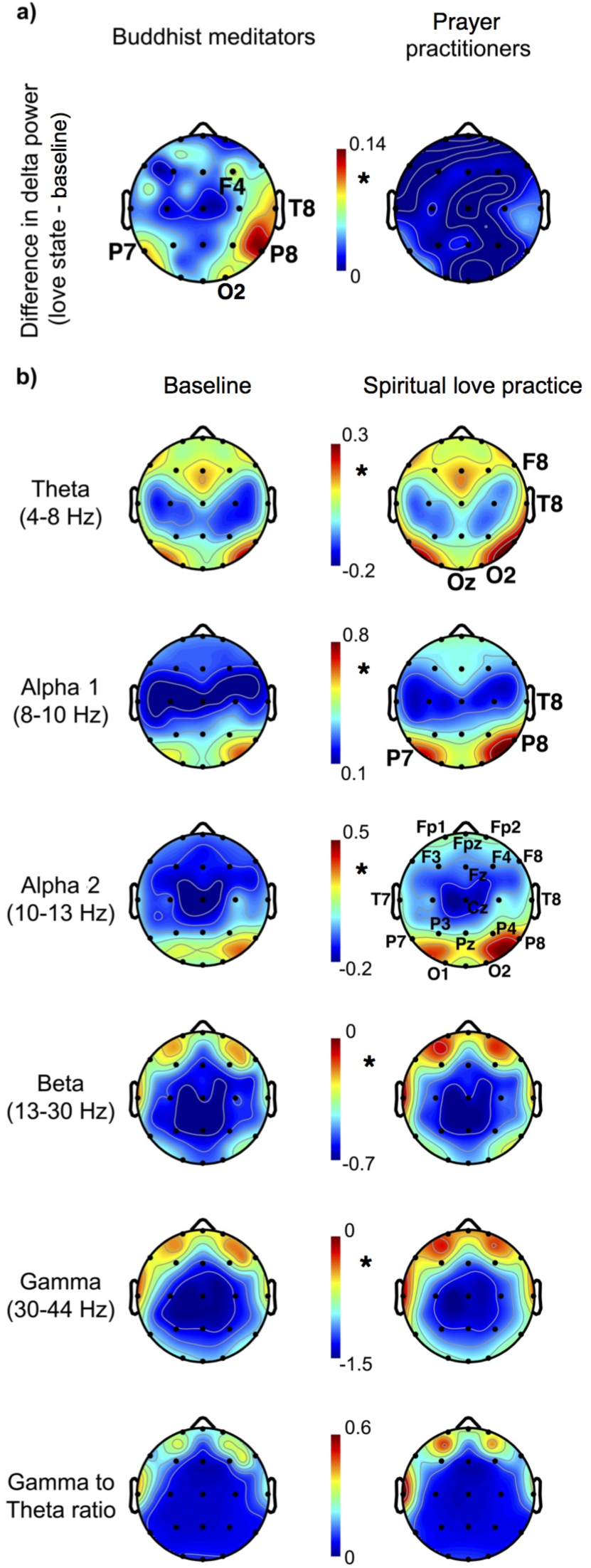
The neural correlates of both forms of spiritual love relative to baseline activity. a) A topographical map showing an increase in delta power during the LKM practice in Buddhist meditators (left panel) but not during prayer (right panel) when comparing the love state to baseline. b) Topographical maps for theta, alpha 1, alpha 2, beta, gamma and the gamma to theta ratio are shown on the left for the baseline condition and on the right for the spiritual love practice. Main effects are reported for the beta and gamma bands, showing increased power during the love practices. Interactions effects, when followed up, showed significant increase of theta, alpha 1 and alpha 2 power during the spiritual love practices compared to baseline. There was a trend towards an increase in the ratio of gamma to theta power, however this failed to reach statistical significance.

Significant increases in the power of theta, alpha 1, alpha 2, beta and gamma frequency bands were observed when averaged across both the spiritual love practices and compared to baseline. A significant two way interaction between state and electrode was observed in the theta frequency band (*F*(20, 500) = 2.522, *p* = 0.036, 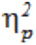 = 0.092, ε = 0.236; Figure 4b). Pairwise comparisons revealed that the increased theta power during the love state relative to baseline occurred at F8, T8, Oz and O2 (*p*<0.05 for all electrodes). Similarly, a significant interaction between state and electrode was observed in the alpha 1 frequency band (*F(*20, 500) = 4.686, *p* <0.001, 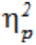 = 0.158, ε = 0.290; Figure 4b) with pairwise comparisons revealing this effect at electrodes T8, P7 and P8 (*p*< 0.05). For the alpha 2 frequency, a significant interaction between state and electrode was present (*F*(20, 500) = 2.729, *p* = 0.010, 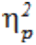 = 0.098, ε = 0.349); Figure 4b), and pairwise comparisons showed global increases in alpha 2 power at electrodes across the brain during the love state relative to baseline at Fpz, Fp1, Fp2, Fz, F3, F4, F8, Cz, T7, T8, Pz, P3, P4, P7, P8, O1 and O2 (*p*<0.05 for all electrodes). For the beta frequency band, a main effect of state revealed a significant increase during the spiritual love practices compared to baseline (*F*(1, 25) = 8.425, *p* = 0.008, 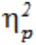 = 0.252; Figure 4b). Similarly, a main effect of state showed a significant increase in gamma power during the love practices relative to baseline (*F*(1, 25) = 6.704, *p* = 0.016, 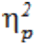 = 0.211; Figure 4b). Lastly, the main effect of state revealed a trend level increase in gamma to theta power during the love state relative to baseline (*F*(1, 25) = 3.965, *p* = 0.057, 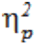 = 0.252; Figure 4b). No interaction or main effects were observed in the ratio of the gamma frequency to the alpha 1 (*F(*20, 500) = 0.806, *p* = 0.486, 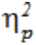 = 0.031, ε = 0.138) and alpha 2 frequency bands (*F*(20, 500) = 0.356, *p* = 0.846, 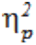 = 0.014, ε = 0.208).

## Discussion

The current study examined the effects of two different types of spiritual practices that cultivate love, a Buddhist loving kindness meditation and a Christian-based prayer, which involves participants generating a love for God and asking to receive love from God. While a large number of studies have examined the effects of meditation on the brain (Cahn & Polich, 2006), this is only the second study to examine the neural correlates of the LKM using EEG. In the first study, highly experienced Tibetan Buddhist monks who had between 10,000 to 50,000 hours of meditation practice were examined (Lutz *et al*., 2004). In contrast, the current study examined relatively inexperienced meditators, with only 800-3800 hours of practice. This allowed us to directly compare the aforementioned group of meditators with a group of similarly experienced practitioners of a Christian-based faith, in terms of hours of practice, who were also matched in age and ethnic background.

Using EEG to measure spectral changes in neural activity, we found that the LKM practice generated significant increases in the delta, alpha 1, alpha 2 and beta frequency power compared to baseline. When comparing prayer to baseline, significant increases were observed in the alpha 1 and gamma frequency bands, with increases in the alpha 2 and beta frequency power almost reaching significance. In addition, an increase in the ratio of gamma to theta frequencies was observed during prayer compared to baseline. Comparing the two spiritual love states, analysis revealed a significant increase in the delta frequency band only during the LKM practice, whereas increases in theta, alpha 1, alpha 2, beta and gamma frequency bands were common to both forms of love.

### Neural correlates of the LKM

Significant increases in the delta frequency band over right frontal, right temporal, left and right parietal and right occipital electrodes were observed when comparing the LKM practice to rest and when comparing the LKM practice to prayer. In a previous study where EEG patterns were compared during the presentation of pictures of faces of loved ones and pictures of “unappreciated” or “unknown" people, widespread increases in delta band activity across the brain were found only during the presentation of faces of loved ones (Basar *et al*., 2008). The LKM practice involves visualising a loved one, experiencing love for that person, and then directing that feeling of love to other people, with the eventual aim of generating a state of unconditional loving-kindness and compassion for all people. Thus the widespread increase in delta power that we observed during the LKM practice in the current study may reflect activation of neural regions involved in generating love for another person, which is a core preliminary requirement of LKM. The primary function of delta rhythms is associated with sleep and drowsiness, however in awake individuals delta activity has been correlated with inhibitory functions (Niedermeyer & Lopez da Silva, 1993). Increased delta power in the right frontal region is suggestive of selective inhibition of this brain region, and is consistent with reports that activation of the left prefrontal cortex compared to the right prefrontal cortex is associated with positive affect and approach behaviours (Coan & Allen, 2004; Davidson *et al*., 2004).

An increase in alpha 1 band power over a right temporal electrode (T8) and a right parietal electrode (P8) was also observed during the LKM practice, while an increase in alpha 2 power was seen averaged across all electrodes. Over the past few decades increases in alpha power have been widely reported during meditation practices, and have been associated with relaxation, lower anxiety and more positive affect (Aftanas & Golocheikine, 2002; Cahn & Polich, 2006; Lagopoulos *et al*., 2009). Alpha 1 amplitude, in particular, has been associated with elevated mood (Petsche *et al*., 1997) and is an index of internalised attention, alertness and expectancy (Klimesch *et al*., 1998; Aftanas & Golocheikine, 2001), while improved memory performance has been correlated with higher alpha 2 power (Klimesch *et al*., 2006), and alpha 2 levels have been suggested to be an indicator of the engagement of brain regions in task performance (Travis & Shear, 2010). The increased alpha power that is observed over posterior and central regions in this study may be reflective of the increased working memory load and attentional processes (Jensen *et al*., 2002) required for performing the LKM practice compared to baseline. In clinical studies, increased activity in right temporal and right parietal areas has been associated with increased anxiety (Mathersul *et al*., 2008). Thus the increase in alpha 1 power in the right temporal and right parietal region observed in this current study suggests that the lower anxiety levels generated through the LKM practice may result from inhibition of these regions, since alpha oscillations are known to play a role in cortical tuning through inhibition (Ergenoglu *et al*., 2004; Klimesch *et al*., 2007).

Increased beta power was also observed during the LKM practice when averaged across all electrodes. Both beta 1 and beta 2 activity have been correlated with executive function and attentional processes (Razumnikova, 2007; Travis & Shear, 2010), and beta 2 has been specifically associated with creative thinking (Razumnikova, 2007). Since the mediators used in the current study were only moderately experienced, sustained concentration is required to maintain the affective state generated through the LKM practice. Thus the increase in alpha 2 and beta activity observed in the current study may reflect the mental effort of sustaining the meditation practice and the generation of a feeling of love. In a previous study examining the LKM in experienced meditators, increases in the gamma frequency band during the practice were reported that correlated with the level of experience of the participants (Lutz *et al*., 2004). We found no increase in the gamma frequency band during the LKM compared to baseline in our moderately experienced meditators, which may be attributable to the relative inexperience of the meditators used in our study.

### Neural correlates of prayer

The spectral changes in EEG patterns observed during prayer showed widespread increases in the alpha 1 frequency band, while increased gamma frequency activity was observed when averaged across all electrodes, as was an increase in the gamma to theta power ratio. A trend towards an increase was observed in the alpha 2 and beta frequency bands, however this did not reach statistical significance.

During prayer, the increase in alpha 1 activity was observed at multiple sites across frontal, central, temporal, parietal and occipital regions. Since activity in the alpha 1 frequency band has been associated with improved mood (Petsche *et al*., 1997), the widespread increase in alpha 1 activity may reflect the enhanced positive affect associated with the experience of this type of spiritual love. The participants who carried out prayer report that it does not require a mental effort to sustain a state of prayer, whereas the LKM practice involves an ongoing concentrative process in meditators of the level of experience used in our study. Thus the different activation patterns of alpha 1, alpha 2 and beta frequency bands seen when comparing the two spiritual love practices to rest may be due to differences in attention and concentration levels required to perform the two practices.

Activity in the gamma frequency band reflects local neuronal processing, and is proposed to be the frequency responsible for the binding of neural activity of large areas of the brain to create a unified conscious experience (Joliot *et al*., 1994; Singer, 1999; von Stein & Sarnthein, 2000). High frequency oscillations in the gamma frequency range can be contaminated by eye saccades and muscle tone (Goncharova *et al*., 2003; Whitham *et al*., 2007), and therefore results showing increased gamma power should be interpreted with caution. However the current study showed a generalized increase in gamma frequency during the prayer compared to rest averaged over all electrodes, and not specifically at frontal electrodes, which would be more easily affected by muscle tone. This increase in gamma activity may reflect the coordination of activity across brain regions, forming a unified mental construction of the experience of this form of spiritual love. In particular, an increase in gamma frequency over prefrontal cortex has been shown to be associated with feelings of happiness and love (Rubik, 2011). The ratio of gamma to theta activity was also significantly increased during prayer. The slower activity in the alpha and theta frequency range has been proposed to be important for coordinating activity over more spatially separate regions of the brain, allowing the integration of distinct neural networks with differing functions (von Stein & Sarnthein, 2000). Specifically gamma and theta activity have been closely linked together as arising from the neural basis of short-term or working memory (Ward, 2003).

The only other previous study to examine neural activity with EEG during a mystical experience, i.e. an experience where participants reported a close connection with God, was carried out in Carmelite nuns (Beauregard & Paquette, 2008). The results of that study showed an increase in gamma activity over right temporal and parietal regions, consistent with increase in gamma power observed in the current findings, and it also reported increases in theta power over frontal, central and parietal regions (Beauregard & Paquette, 2008). In the previous study, participants initiated the mystical state by recalling times when they had felt a union with God, which then reportedly generated a real time mystical experience. Therefore the difference in theta activation seen in the previous study compared to the current study may be due in part to activation of memory systems during the recollection, since theta oscillations are known to play a role in memory (Rutishauser *et al*., 2010). In addition, the nuns who attained the mystical state in the previous study may have experienced a more intense blissful state, which has been associated with enhanced frontal theta (Aftanas & Golocheikine, 2001), than the prayer state examined in the current set of moderately experienced participants. Future experiments directly examining the difference between moderately experienced and highly experienced Christian participants engaging in prayer, together with a comparison memory task, would be required to resolve this.

### Comparison of the two spiritual love practices

Combined examination of the LKM and prayer compared to rest revealed significantly increased theta power in right frontal and temporal regions and in right and central occipital regions; increased alpha 1 at a right temporal electrode and bilaterally in parietal regions; global increases in alpha 2 power; and significant increases in beta and gamma power averaged across all electrodes. It is notable that an increase in alpha 1 power was observed at electrodes T8 and P8 for the both the LKM and prayer versus baseline activity. In addition to the correlation between right temporal and parietal activity with anxiety as mentioned above (Mathersul *et al*., 2008), an increase in alpha power over the right parietal lobe has also been observed during responses to both positive and negative emotional cues, including happy, sad, afraid, angry, surprised and disgust emotional responses (Yuvaraj *et al*., 2014), consistent with the notion that activity in this region is associated with emotion processing and arousal, rather than the valence of emotion experienced (Heller, 1993).

Increased theta power has been reported during many forms of concentrative meditation (Cahn & Polich, 2006). An increase in frontal theta has also been correlated with enhanced mindfulness, or self-awareness, and with subjective reports of the attainment of a blissful state (Aftanas & Golocheikine, 2001). The neural source of this frontal theta is widely reported to be the anterior cingulate cortex, which is involved in cognitive and affective mechanisms (Asada *et al*., 1999) (Ishii *et al*., 1999). Activity in the anterior cingulate cortex is negatively correlated with sympathetic activity (Kubota *et al*., 2001) and positively correlated with parasympathetic activity (Tang *et al*., 2009), consistent with physiological activity associated with lessened anxiety experienced during love compared to baseline.

Increased theta activity is also correlated with the ability to encode new information (Klimesch, 1999) and the formation of new memories (Rutishauser *et al*., 2010). At the cellular level, theta oscillations and high frequency activation in the gamma frequency range can induce synaptic plasticity (Larson *et al*., 1986; Buzsaki & Draguhn, 2004). Thus the significant increases in theta activity and absolute gamma power, together with the trend towards an increase in the gamma to theta power ratio during the two spiritual love practices, in particular over frontal and temporal regions, is suggestive that these practices may lead to the formation of new memories via the alteration of neuronal circuitry.

Comparison of prayer with resting state showed widespread changes in neural activity across the brain, across a range of frequency bands. These changes showed similarities with the LKM, but also a difference in the delta frequency band. It is the changes that were observed during both prayer and the LKM compared to baseline; the increase in activity in the theta, alpha 1, alpha 2, beta and gamma frequency bands, that are likely to reflect the neural correlates of the experience of spiritual love. In contrast the increase in delta activity in the brain during the LKM may be a feature of the meditative practice itself as opposed to spiritual love. Therefore this study suggests that despite the marked differences in technique and religious orientation of the Buddhist participants and participants who engaged in prayer, i.e. non-theistic versus theistic respectively, there appear to be substantial similarities in the neural correlates of spiritual love.

Mindfulness practices, which are implicit in the LKM practice, are effective in the clinical treatment of anxiety disorders and depression (Baer, 2003; Grossman *et al*., 2004; Sundquist *et al*., 2014). The increase in theta and gamma frequencies observed in both the relatively inexperienced Buddhist meditators and participants carrying out prayer suggest that practicing these forms of spirituality over a relatively short time period may be a useful approach for improving positive affect in the long-term by altering the neural circuitry of the brain (Esch & Stefano, 2005; Fell *et al*., 2010). In addition, positive affect has been shown to be beneficial to overall physical health, by boosting immune system function and reducing stress, which is linked to a number of chronic diseases such as cardiovascular disease and cancer (Kiecolt-Glaser *et al*., 2002), suggesting that both practices that cultivate love may be of assistance to physical as well as mental health.

## Acknowledgements

This research was funded by the National Health and Medical Research Council of Australia. I’d like to thank Professor Perry Bartlett for supporting this study at the Queensland Brain Institute, Professor Jason Mattingley for advice on the design of the experiments, and Oscar Jacoby for programming the software for the experimental design. I would also like to acknowledge and thank other collaborators at the Queensland Brain Institute involved in the project. The author has no competing interests.

